# Alteration of gut microbiota with a broad-spectrum antibiotic does not impair maternal care in the European earwig

**DOI:** 10.1101/2020.10.08.331363

**Authors:** Sophie Van Meyel, Séverine Devers, Simon Dupont, Franck Dedeine, Joël Meunier

## Abstract

The microbes residing within the gut of an animal host often increase their own fitness by modifying their host’s physiological, reproductive, and behavioural functions. Whereas recent studies suggest that they may also shape host sociality and therefore have critical effects on animal social evolution, the impact of the gut microbiota on maternal care remains unexplored. This is surprising, as this behaviour is widespread among animals, often determines the fitness of both juveniles and parents, and is essential in the evolution of complex animal societies. Here, we tested whether life-long alterations of the gut microbiota with rifampicin - a broad-spectrum antibiotic - impair pre- and post-hatching maternal care in the European earwig. Our results first confirm that rifampicin altered the mothers’ gut microbial communities and indicate that the composition of the gut microbiota differs before and after egg care. Contrary to our predictions, however, the rifampicin-induced alterations of the gut microbiota did not modify pre- or post-hatching care. Independent of maternal care, rifampicin increased the females’ feces production and resulted in lighter eggs and juveniles. By contrast, rifampicin altered none of the other 21 physiological, reproductive and longevity traits measured over the 300 days of a female’s lifetime. Overall, these findings reveal that altering the gut microbiota with a large spectrum antibiotic such as rifampicin does not necessarily affect host sociality. They also emphasize that not all animals have evolved a co-dependence with their microbiota and call for caution when generalizing the central role of gut microbes in host biology.

## Introduction

Almost all animals harbour a gut microbiota, i.e. a community of microorganisms residing within the gut of the host [1]. Some of these gut microbes have long been known for their pathogenic effects in the hosts [2] and others for the benefits they provide to the hosts in terms of nutritional mutualism [3]. Over the last decades, however, a growing number of studies has been revealing that the effects of gut microbes are much more diverse than previously thought and shape numerous physiological, reproductive, and behavioural functions of the host [4]. In the fruit fly Drosophila melanogaster, for instance, the gut microbiota is associated with hormone signalling, metabolism and ageing [5]. Gut microbes can also shape the hosts’ immunocompetence and resistance against pesticides and toxic plant metabolites [6], such as in the mosquito Anopheles stephensi [7], the bean bug Riptortus pedestri [8] and the wasp Nasonia vitripennis [9]. Similarly, the gut microbiota is a key parameter in host reproduction and mating incompatibilities, as found in the fruit fly Bactrocera minax [10], the terrestrial isopod Armadillidium vulgare [11], and the parasitic wasp Asobara tabida [12]. Finally, gut microbes shape the expression of numerous host behaviours, such as offspring activity in the stinkbug Megacopta punctissima [13], and different tasks in honeybees [14].

Recent studies also suggest that gut microbiota can play a critical role in the sociality of their hosts by shaping the expression and nature of social interactions and/or by transforming mediators of social aggregation. For instance, family-living rats with a diet-altered gut microbiota exhibit deficient sociability and increased social avoidance [15,16]. Antibiotic-induced modifications of gut microbiota also alter the chemical signatures of social hosts and lead to higher levels of aggressiveness toward conspecifics in the leaf-cutting ant Acromyrmex echinator [17] and the honeybee Apis mellifera [18]. Finally, alteration of the gut microbiota reduces the production of aggregation pheromones in the swarm-living desert locust Schistocerca gregaria [19] and diminishes the presence of aggregation pheromones in feces of the gregarious German cockroach Blattella germanica [20].

Despite these causal and correlative links between the hosts’ gut microbial communities and sociality, the role of gut microbes on the expression of parental care remains experimentally unexplored. This is surprising, because this form of social behaviour is present in a large and taxonomically diverse number of animal species [21], has considerable effects on the fitness of both juveniles and parents [22] and because shedding light on this link may provide crucial information on the role of gut microbes in the early evolution of complex animal societies [23]. On one hand, gut microbes could indirectly alter the investment in parental care, because parents are expected to adjust their level of care to their own condition [24] and altered gut microbial communities can lower these conditions in multiple ways (see above). On the other hand, gut microbes could serve as a direct promoter of parental care because, by enforcing the expression of care in adult hosts, parental gut microbes could maximize their chances to reach novel hosts [25–30](but see [31]). The transfer of gut microbes through parental care has been reported in several insect species, such as the stinkbug *Parastrachia japonensis* [32], the Japanese common plataspid stinkbug *Megacopta punctatissima* [33], the wood cockroach *Cryptocercus punctulatus* [34] and the wood-feeding termite *Reticulitermes grassei* [35]. However, we still know very little about whether and how the gut microbiota influences the investment in parental care.

In this study, we address this gap in knowledge by investigating whether gut microbiota alteration with rifampicin - a broad-spectrum antibiotic - impaired the expression of pre- and post-hatching maternal care in the European earwig *Forficula auricularia*. In this omnivorous insect, mothers tend their first clutch of eggs over winter and their second and terminal clutch (when present) during spring. During these periods, mothers stop their foraging activity and provide egg care in the forms of protection against desiccation, predators and pathogens [36]. Egg care also mediates the transfer of microbes with antifungal properties to eggshell in the maritime earwig *Anisolabis maritima* [37], a process that possibly occurs in the European earwig [38]. Upon egg hatching, *F. auricularia* mothers continue tending their brood of newly emerged juveniles (called nymphs) for two weeks, during which they provide care in the forms of fierce protections against predators, grooming behaviours, and food provisioning through regurgitation [39]. Pre-hatching care is necessary to ensure egg development and hatching [38], whereas post-hatching care is facultative for the development and survival of nymphs [40]. Earwig females present important inter-individual variation in the expression of maternal care within populations [41,42], and this variation is partly inherited from the parents [43] and partly depends on environmental inputs, such as the social environment or food resources [44–46]. To the best of our knowledge, this is the first study to explore the microbiome composition of the European earwig.

We altered the gut microbiota of *F. auricularia* females by feeding them with rifampicin during their entire adult lifetime (about 14 months) and measured whether and how it affected gut microbial communities, maternal care, and other life-history traits. Specifically, we first determined how the antibiotic treatment alters the diversity and structure of the gut bacterial community of females at two periods of their life-cycle (just before the production and at the hatching of their 1st clutch eggs) by sequencing 16S rRNA gene (V3-V4 region) amplicons. We then tested the effects of rifampicin on the expression of four pre- and two post-hatching forms of maternal care toward 1st clutch eggs and nymphs, respectively. Finally, to disentangle whether the potential link between gut microbiota alteration and the level of maternal care is direct and/or indirect, we investigated the effects of rifampicin on 24 other traits measured throughout the mothers’ lifetime and reflecting their general physiological state, investment in future reproduction and longevity.

## Methods

### Insect rearing and rifampicin treatment

The experiment involved a total of 296 *Forficula auricularia* L. (clade B [47]) males and females. These were the first laboratory-born descendants of 74 females field-sampled in Pont-de-Ruan, France, in 2017 and then maintained under standard laboratory conditions [42]. The entire experiment consisted of feeding these 296 earwigs with either rifampicin or water from adult emergence to death, and measuring the effects on mothers’ behaviour, physiology, reproduction, and longevity (Figure S1). To obtain rifampicin- and control-treated mothers, we first isolated two virgin males and two virgin females per family (n = 74 families) four days after adult emergence, and then fed them for two weeks with green-coloured pollen pellets mixed with either 10 μL of rifampicin (Sigma-Aldrich, PHR1806; 0.2 mg/ml) or 10 μL of water. During these two weeks, each individual was isolated in a Petri Dish grounded with a humid filter paper that was changed twice a week (together with the treated food) to limit the risk of self-transplantation of gut microbes through the consumption of feces deposited on paper. The production of green-coloured feces was always observed, confirming the consumption of rifampicin-treated food. At the end of these two weeks, we used the 296 earwigs to set up 148 mating pairs composed of 1 virgin female and 1 virgin male from the same family and the same treatment. There are only limited signs of inbreeding depression in this species [48] and sib-mating allowed us reducing the risk of poor reproductive success due to possible inter-familial cytoplasmic incompatibility between certain bacterial strains, as reported with Wolbachia and Cardinium in several arthropod species [49,50]. Each of the resulting 148 mating pair received a standard food source mostly composed of agar, carrots, pollen, and cat and bird dry food [42] twice a week for two months. This food was mixed with either 10 μL of rifampicin (0.2 mg/ml) or 10 μL of water to follow-up on the previous treatments. After these two months, females were isolated and maintained under winter conditions to mimic natural dispersal, allow oviposition, and subsequently measure four forms of egg care (details below) [42]. Mothers were not provided with food and thus not treated with rifampicin from oviposition to egg hatching, as mothers typically stop foraging during this period [40]. One day after egg hatching, each family was maintained under spring conditions and fed with the standard food source mixed with either 10 μL of rifampicin (0.2 mg/ml) or 10 μL of water according to the pre-oviposition treatment. The food and treatment were renewed twice a week. We measured three forms of maternal care towards juveniles during the following 14 days (details below), which corresponds to the duration of family life in this species [42]. Nymphs were then discarded from the experiment to allow newly isolated mothers to produce a 2nd clutch. These mothers were then maintained under spring conditions and continued to receive the same treatment (rifampicin or water) until they die. Except when stated otherwise, individuals were always maintained in Petri dishes (diameter 9cm) lined with non-sterile moistened sand.

Rifampicin is considered one of the most potent and broad-spectrum antibiotics due to its high-affinity binding to the RNAP β subunit, which causes the inhibition of the bacterial DNA-dependent RNA polymerase RNAP by directly blocking the RNA elongation path [51]. It is also commonly used to experimentally alter gut microbial communities in insects (e.g. [52–54]). The high dose of rifampicin used in this study (about 10 times higher than the dose generally used in smaller insect species [53,54]) was chosen to ensure gut microbial alteration and because it did not trigger an excess of mortality in the German cockroach [52], an insect that is about the size of the European earwig.

### Effects of rifampicin on the gut microbiota

To test whether and how rifampicin treatment altered the earwigs’ gut microbial communities, we extracted the gut of 10 females per treatment (n total = 20) on the day we observed the first oviposition of their 1st clutch (i.e. about 2 months after being fed with or without rifampicin), and 10 rifampicin- and 8 water-treated females one day after their 1st clutch eggs have hatched (i.e. about 1 month later; Figure S1). For gut extraction, we first anaesthetized each female for 2 min at −20°C and then dissected them in a watch glass with sterilized double Q water. All dissections and manipulations were conducted on a sterilized bench, under a Bunsen burner’s sterility area and using sterile material. Whole individual guts were extracted, placed in 100 μl of T1 extraction buffer (Nucleo-Spin Tissue, Macherey-Nagel), and stored at −80°C until DNA extraction. Two PCR amplifications were performed for each sample in a final volume of 35 μl to amplify a 450-bp portion of the V3–V4 region of the 16S rRNA gene (forward primer: 5′-CTT TCC CTA CAC GAC GCT CTT CCG ATC TAC GGR AGG CAG CAG-3′; reverse primer: 5′-GGA GTT CAG ACG TGT GCT CTT CCG ATC TTA CCA GGG TAT CTA ATC-3′; the Illumina adapters and primers per se are indicated in non-bold and bold, respectively). 50 μl of PCR product were then sent to the GeT-PlaGe genomic platform (GenoToul, Toulouse, France), which performed library preparation and 2 × 250 paired-end Illumina Miseq sequencing according to the manufacturer’s instructions. DNA extractions protocols, sequencing process and bioinformatic pipelines are detailed in the supplementary material.

### Measurements of pre- and post-hatching maternal care

We first measured the effects of rifampicin on four classical forms of earwig maternal egg care: egg grooming, egg defence, delay of maternal return and exploration rate while searching for its eggs [55,56]. Egg grooming, which is used by earwig females to deposit chemical substances on the eggs and to clean eggshell from dirt and fungi [38], was measured 15 days after egg production. We first isolated each mother for 30 min, then returned them to their Petri dish and gently deposited them at a distance of 5 cm from their clutch, and finally video-recorded their behaviours for the subsequent 15 minutes (SONY© Handycam HDR-CX700 camera). The resulting movies were analysed using the software BORIS v4.0.3 [57] and the total duration of egg grooming was defined as the total amount of time each female spent on cleaning eggs with their mandibles [38]. Clutch defence, which reflects the females’ willingness to protect their eggs from a predator attack [44], was measured 16 days after egg production. We standardly poked each female on the pronotum with a glass capillary (one poke per second) and then recorded the number of pokes required until the female moved more than one body length away from the clutch. The delay of maternal return after clutch abandonment [56], which represents the delay after which females return to their clutch after being chased away by a simulated predator attack [44], was measured by recording the time the female took to return to its clutch just after the end of the clutch defence measurement. Finally, the exploration rate, which represents the level of exploration of a novel area by a mother looking for her eggs, was measured 21 days after egg production. We removed each mother from its clutch of eggs, subsequently deposited her at the centre of a square arena (W: 9 cm; L: 9 cm; H: 0.5 cm) covered by a glass sheet, and then video-tracked its activity during 35 min. The video was done under infra-red light, while the individual video tracking and calculation of exploration rate were conducted using the software ToxTrac v2.83 [58].

We then measured the effects of rifampicin on two classical forms of post-hatching maternal care: nymphs defence and exploration rate while searching for its nymphs [44,55]. These two forms of care were measured 10 and 12 days after egg hatching, respectively, following the above-detailed protocols for egg defence and egg searching.

All the measurements of pre- and post-hatching maternal care were conducted in the afternoon and under a red light as earwigs are nocturnal. These measurements were conducted blindly regarding the treatments (rifampicin versus control). The number of replicates for each of our measurements ranged from 9 to 59 per treatment (median = 36 per treatment; details in Tables 1 and S1). The recorded range of values of maternal care is comparable to the range of values obtained in previous studies conducted in other populations [41,42,44] and thus likely reflects the natural variation in maternal care exhibited by earwig females.

**Table 1.**
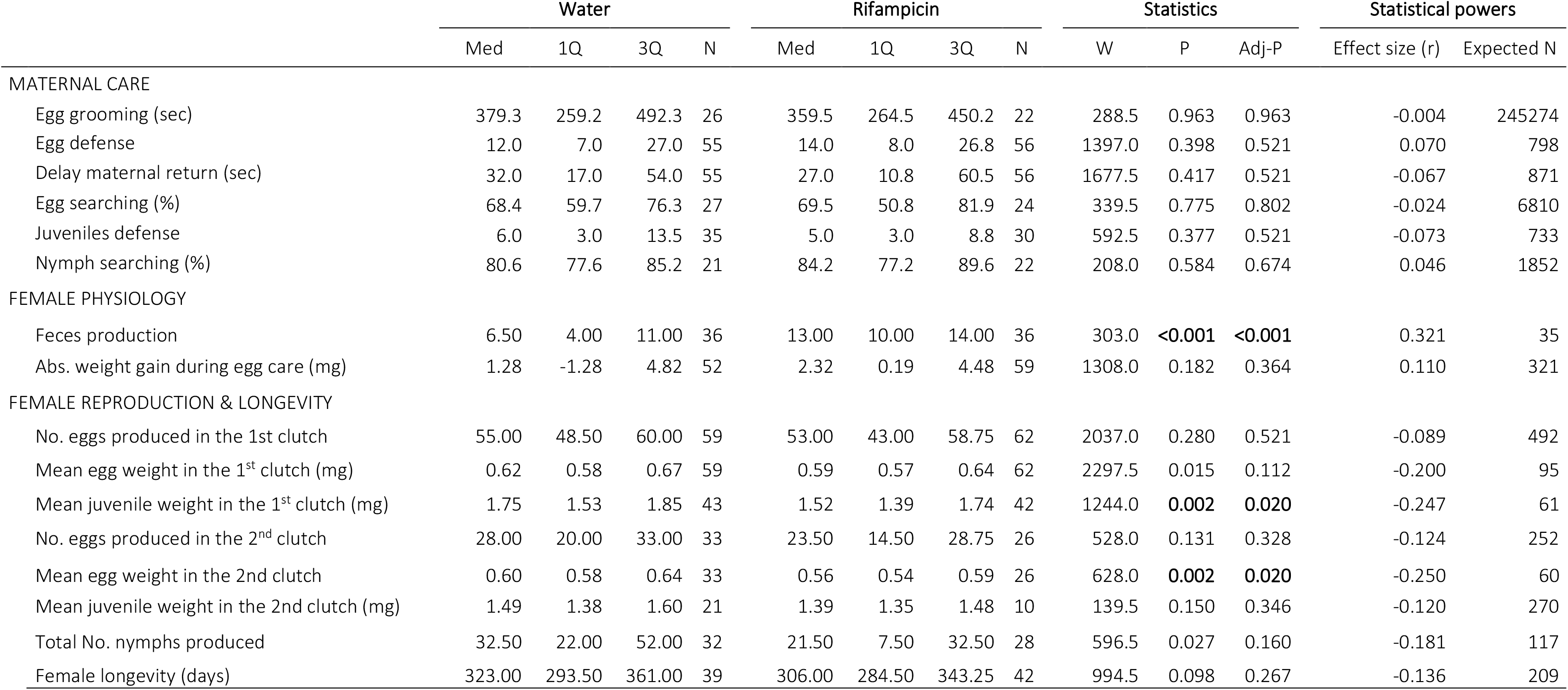
Effects of rifampicin on a representative selection of 16 of the 30 measured traits reflecting maternal care, physiology, reproduction, and longevity. The effects on the 30 traits are presented in table S1. P-values significant after correction for multiple comparisons (Adj-P) are in bold. Med = Median; 1Q and 3Q = first and third quartile, respectively. N = sample size. Expected N = number of replicates per treatment that would have been necessary to obtain a statistically significant difference with a power of 0.8.

|  | Water |  |  |  | Rifampicin |  |  |  | Statistics |  |  | Statistical powers |  |
| --- | --- | --- | --- | --- | --- | --- | --- | --- | --- | --- | --- | --- | --- |
|  | Med | 1Q | 3Q | N | Med | 1Q | 3Q | N | W | P | Adj-P | Effect size (r) | Expected N |
| MATERNAL CARE |  |  |  |  |  |  |  |  |  |  |  |  |  |
| Egg grooming (sec) | 379.3 | 259.2 | 492.3 | 26 | 359.5 | 264.5 | 450.2 | 22 | 288.5 | 0.963 | 0.963 | -0.004 | 245274 |
| Egg defense | 12.0 | 7.0 | 27.0 | 55 | 14.0 | 8.0 | 26.8 | 56 | 1397.0 | 0.398 | 0.521 | 0.070 | 798 |
| Delay maternal return (sec) | 32.0 | 17.0 | 54.0 | 55 | 27.0 | 10.8 | 60.5 | 56 | 1677.5 | 0.417 | 0.521 | -0.067 | 871 |
| Egg searching (%) | 68.4 | 59.7 | 76.3 | 27 | 69.5 | 50.8 | 81.9 | 24 | 339.5 | 0.775 | 0.802 | -0.024 | 6810 |
| Juveniles defense | 6.0 | 3.0 | 13.5 | 35 | 5.0 | 3.0 | 8.8 | 30 | 592.5 | 0.377 | 0.521 | -0.073 | 733 |
| Nymph searching (%) | 80.6 | 77.6 | 85.2 | 21 | 84.2 | 77.2 | 89.6 | 22 | 208.0 | 0.584 | 0.674 | 0.046 | 1852 |
| FEMALE PHYSIOLOGY |  |  |  |  |  |  |  |  |  |  |  |  |  |
| Feces production | 6.50 | 4.00 | 11.00 | 36 | 13.00 | 10.00 | 14.00 | 36 | 303.0 | **<0.001** | **<0.001** | 0.321 | 35 |
| Abs. weight gain during egg care (mg) | 1.28 | -1.28 | 4.82 | 52 | 2.32 | 0.19 | 4.48 | 59 | 1308.0 | 0.182 | 0.364 | 0.110 | 321 |
| FEMALE REPRODUCTION & LONGEVITY |  |  |  |  |  |  |  |  |  |  |  |  |  |
| No. eggs produced in the 1<sup>st</sup> clutch | 55.00 | 48.50 | 60.00 | 59 | 53.00 | 43.00 | 58.75 | 62 | 2037.0 | 0.280 | 0.521 | -0.089 | 492 |
| Mean egg weight in the 1<sup>st</sup> clutch (mg) | 0.62 | 0.58 | 0.67 | 59 | 0.59 | 0.57 | 0.64 | 62 | 2297.5 | 0.015 | 0.112 | -0.200 | 95 |
| Mean juvenile weight in the 1<sup>st</sup> clutch (mg) | 1.75 | 1.53 | 1.85 | 43 | 1.52 | 1.39 | 1.74 | 42 | 1244.0 | **0.002** | **0.020** | -0.247 | 61 |
| No. eggs produced in the 2<sup>nd</sup> clutch | 28.00 | 20.00 | 33.00 | 33 | 23.50 | 14.50 | 28.75 | 26 | 528.0 | 0.131 | 0.328 | -0.124 | 252 |
| Mean egg weight in the 2nd clutch | 0.60 | 0.58 | 0.64 | 33 | 0.56 | 0.54 | 0.59 | 26 | 628.0 | **0.002** | **0.020** | -0.250 | 60 |
| Mean juvenile weight in the 2nd clutch (mg) | 1.49 | 1.38 | 1.60 | 21 | 1.39 | 1.35 | 1.48 | 10 | 139.5 | 0.150 | 0.346 | -0.120 | 270 |
| Total No. nymphs produced | 32.50 | 22.00 | 52.00 | 32 | 21.50 | 7.50 | 32.50 | 28 | 596.5 | 0.027 | 0.160 | -0.181 | 117 |
| Female longevity (days) | 323.00 | 293.50 | 361.00 | 39 | 306.00 | 284.50 | 343.25 | 42 | 994.5 | 0.098 | 0.267 | -0.136 | 209 |

### Measurements of the 24 other life-history traits in mothers

We used standard protocols to test the effects of rifampicin on 7 proxies of female physiology, as well as 16 proxies of female reproduction and female longevity [42,59]. Proxies of females’ physiology were the number of feces pellets produced per 24 hours (a number positively associated with their digestive/foraging activity [60]) and the gain in fresh weight between two life stages. Feces production was measured two months after the beginning of the treatments. Females were isolated in a new Petri Dish for 24 hours, after which we counted the number of feces pellets present on the ground. The weight gained by each female was measured between the days of adult emergence and oviposition, and between the days of oviposition and egg hatching. Proxies of female reproduction were measured in the 1st and 2nd clutches (if any) by counting the number of eggs produced, the number of days between oviposition and egg hatching (egg development time), and by measuring the mean egg weight at oviposition, the egg hatching rate, and the mean offspring weight at egg hatching. We also counted the number of days between adult emergence and oviposition (days until 1st clutch oviposition), between the females’ isolation after family life and 2nd clutch oviposition (days until 2nd clutch production), and between adult emergence and death (female longevity). We finally assessed whether females produced a 2nd clutch (yes = 1 or no = 0) and females’ reproductive allocation between the two clutches (i.e. the females’ reproductive strategy [42]) defined as the number of 2nd clutch eggs divided by the total number of eggs produced by a female. Overall, weighing was done to the nearest 0.01 mg using a microbalance (OHAUS© Discovery DV215CD). Sample sizes are detailed in Tables 1 and S1.

### Statistical analyses

#### Analyses of the α and β-diversity indices

The structure, composition and diversity of the microbial communities were based on the 161 identified bacterial Operational Taxonomic Units (OTUs) (see results) and analysed using PHYLOSEQ R package [61] implemented in the FROGSSTAT Phyloseq tools [62]. Diversity within the gut microbial communities (alpha-diversity) was assessed using two richness indices, which estimate the number of OTUs in the microbiome with correction for subsampling (Chao1; ACE), and three metrics that aim to measure diversity by accounting for evenness or homogeneity (Shannon, Simpson, Inverse Simpson, and Fisher) [63]. Diversity between the gut microbial communities (beta-diversity) was assessed using 4 measures of community similarity: 1) Jaccard indice, which does not consider phylogeny of OTUs but takes into account their presence/absence; 2) Bray Curtis dissimilarity, which does not consider the phylogeny but considers the number of reads assigned to an OTU (i.e. its abundance); 3) UniFrac indice, which considers phylogeny but not abundance; and finally 4) Weighed UniFrac indice, which considers both phylogeny and abundance. The metrics were analysed individually using either a General Linear Model for α-diversity, or a Permutational Multivariate Analysis of Variance Using Distance Matrices (PERMANOVA) for β-diversity. In these models, the values (or distance matrix for β-diversity) of each index were entered as a response variable, while the treatment (rifampicin or water), the sampling stage of the female (before 1st oviposition or at 1st clutch egg hatching) and the interaction between them were used as fixed factors. When required, a post-hoc analysis was conducted by splitting the data set according to the sampling stage and then conducting PERMANOVA on each of the two resulting subsets. To correct for multiple testing in these post-hoc analyses, the significance level was adjusted to alpha = 0.0375 using the Mean False Discovery Rate approach [64].

#### Analyses of the life-history traits

Although the presented experimental design was originally paired, i.e. 2 females per family distributed among the two treatments, the 38 life-history traits were often measured in only one of the pairs (see Tables 1 and S1). This was mostly due to time constraints, and because some females died during the 18-months course of this experiment. These overall led to critical reductions in the number of replicates that could be involved in a paired statistical approach (details in Table S2). We, therefore, analysed the effects of rifampicin on the 30 measurements using a series of 29 exact Mann Whitney U tests and 1 Pearson’s Chi-squared test (for 2nd clutch production), in which we compared the values of all the available replicates fed with rifampicin to the values of all the available replicates fed with water. Note that the results do not qualitatively change when we use paired analyses with the associated smallest sample sizes (results presented in Table S2). To correct for the inflated risk of Type I errors due to multiple testing, all p-values were adjusted using the False Discovery Rate (FDR) method [64]. To confirm the robustness of non-significant results, we also calculated the effect size r of each analysis and the number of replicates that would have been required to detect a statistically significant effect with this effect size and a statistical power of 0.8. All these analyses were conducted using the software R v4.0.2 (http://www.r-project.org) loaded with the packages exactRankTests, car, rcompanion and pwr.

## Results

### Description of the earwig gut microbiota

A total of 1636435 sequenced reads of the 16S rRNA V3-V4 region were obtained from the 38 female earwig gut samples. After sequence processing, this number went down to 1130241, with 21069 to 35385 sequences per sample (median = 30595.5). The sequences were aggregated and filtered in a total of 161 unique OTUs, which were resolved down to the family or genus level to increase the confidence in the taxonomic assignation. All detailed information on OTUs is given in Table S3. More than 99.90% of the sequences were assigned to four bacterial phyla: Proteobacteria (65.94%), Firmicutes (21.12%), Bacteroidota (9.89%) and Actinobacteriota (2.96%) (Figure 1). The remaining OTUs were assigned to Bdellovibrionota (0.04%) and Patescibacteria (0.04%). The prevalence (i.e. frequency) of these 161 OTUs among the 38 tested females ranged from 0.211 to 1.000 (Table S3). The vast majority of OTUs were found in at least one female in each experimental modality (Table S3), indicating that our rifampicin treatment did not eliminate specific OTUs.

**Figure 1.**
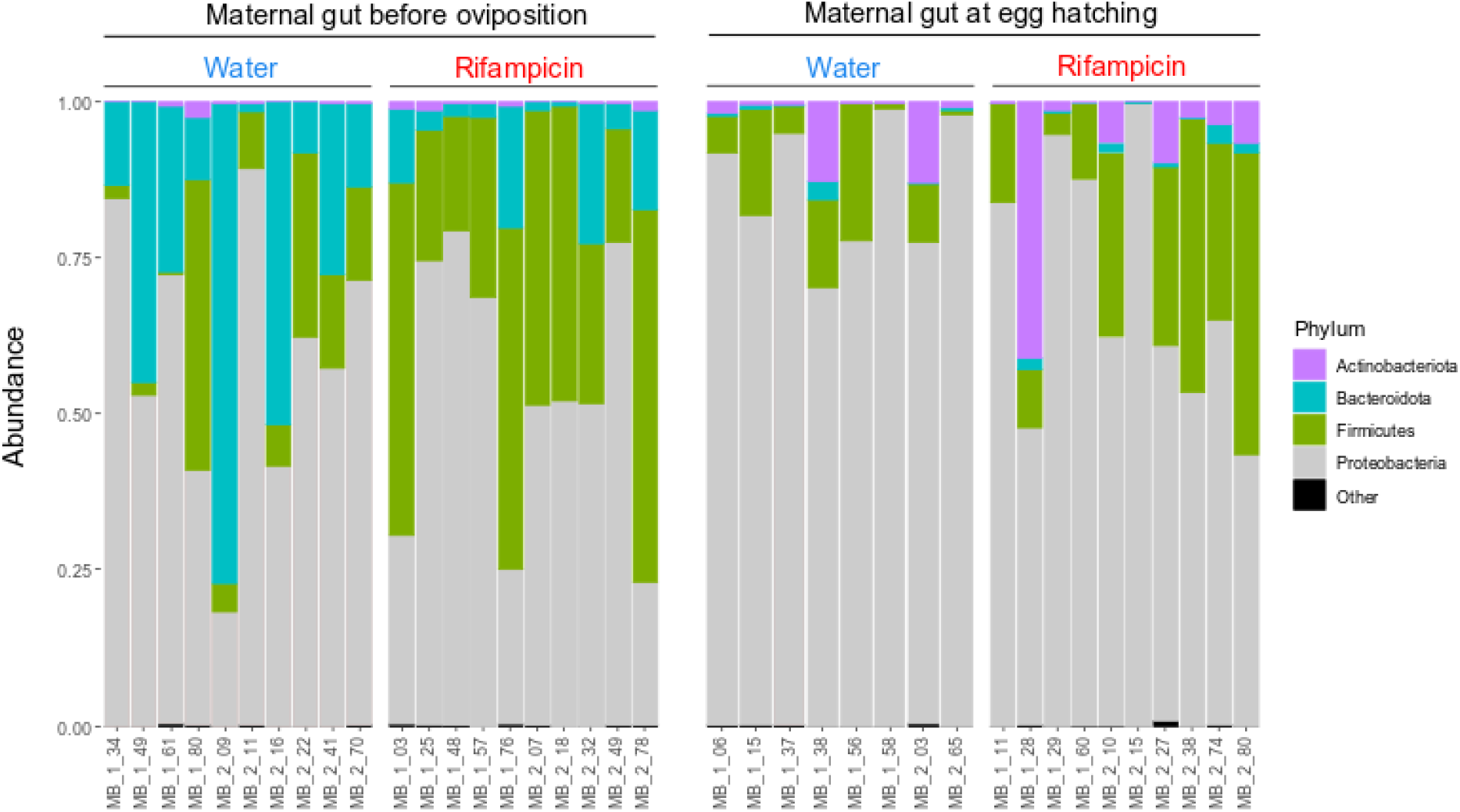
Gut microbial composition in females. Guts were sampled either before oviposition or at the hatching of the 1st clutch in females treated either with water or rifampicin. The ID of each female is provided on the x-axis. These results are presented at the phylum level for clarity, whereas statistical analyses of gut microbial diversity were conducted using OTUs. More details in table S3.

### Comparative analyses of the α and β diversity of the gut microbiota

The gut microbial α-diversity (i.e. species richness) decreased between oviposition and egg hatching when this diversity was measured using Chao1 (F1,34 = 21.63, P < 0.0001), ACE (F1,34 = 24.46, P < 0.0001) and Fisher (F1,34 = 20.85, P < 0.0001; Figure 2) indices. This decrease was, however, not significant when α-diversity was measured using Shannon (F1,34 = 3.18, P = 0.084; Figure 2) and Simpson (F1,34 = 1.60, P = 0.214) indices. Similarly, the α-diversity did not decrease in the rifampicin treatment compared to the control (Chao1: F1,34 = 0.72, P = 0.401; ACE: F1,34 = 0.62, P =0.435; Fisher: F1,34 = 0.59, P = 0.447; Shannon: F1,34 = 1.67, P = 0.205; Simpson: F1,34 = 0.55, P = 0.465; Figure 2), and it was independent of an interaction between female sampling stage and rifampicin treatment (all P > 0.525).

**Figure 2.**
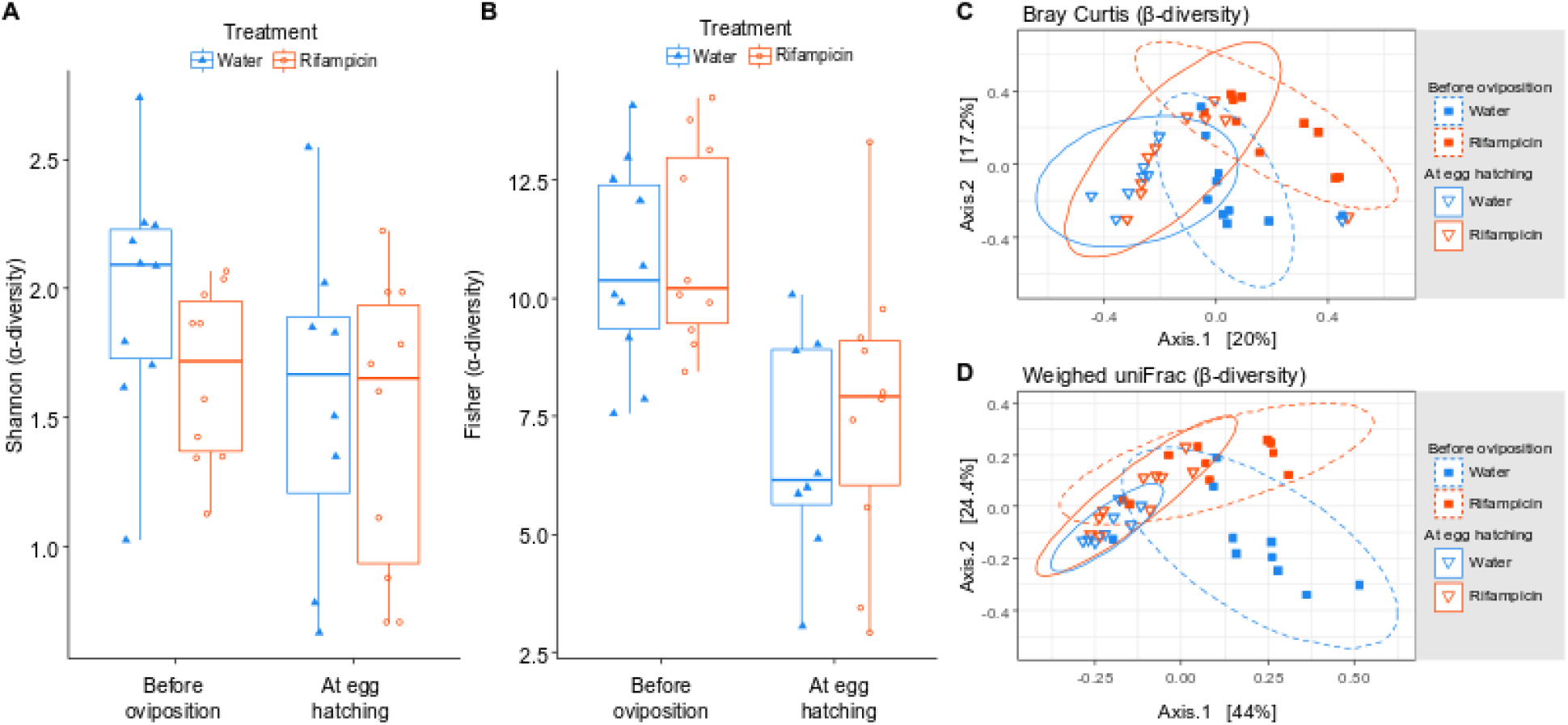
Effects of rifampicin and female sampling stage on gut microbial α- and β-diversities. Guts were sampled either before oviposition or at the hatching of the 1st clutch in females treated either with water or rifampicin. (A, B) Alpha-diversity based on Shannon and Fisher indices as representative of all the tested metrics. Box plots depict median (middle bar) and interquartile range (box), with whiskers extending to 1.5 times the interquartile range and dots/triangles representing values of each sample. (C, D) Beta-diversity based on Bray-Curtis and weighed-uniFrac indices as representative of all the tested metrics. Illustrations report multidimensional scaling (MDS) results, where dots show values and ellipses represent 95% confidence intervals.

The gut microbiota β-diversity (i.e. species composition) overall changed with female sampling stage and rifampicin treatment. This was the case with the four measured indices of β-diversity: Bray-Curtis (Stage: F1,34 = 5.77, P < 0.0001; Rifampicin: F1,34 = 4.23, P < 0.0001), Jaccard (Stage: F1,34 = 7.76, P < 0.0001; Rifampicin: F1,34 = 2.37, P = 0.0036), unweighted UniFrac (Stage: F1,34 = 6.51, P < 0.0001; Rifampicin: F1,34 = 3.39, P = 0.0006) and weighted UniFrac (Stage: F1,34 = 14.10, P < 0.0001; Rifampicin: F1,34 = 6.42, P = 0.0006). In particular, females before oviposition harboured less Actinobacteriota and Proteobacteria compared to females at egg hatching, while rifampicin females overall harboured less Bacteroidota and more Firmicutes compared to untreated females (Figure 1). The interaction between female sampling stage and rifampicin had no effect on the β-diversity measured using all (all P > 0.117) but the weighted UniFrac indices (F1,34 = 2.94, P = 0.026). This interaction reflected an effect of rifampicin on the β-diversity before oviposition (F1,34 = 0.17, P = 0.018) but not at egg hatching (F1,34 = 0.97, P = 0.356).

### Rifampicin and maternal care

We did not detect any effect of rifampicin on the six measured forms of egg and nymph care (Table 1). In particular, rifampicin- and control-fed mothers spent the same amount of time in egg grooming (Figure 3A), showed the same levels of both egg and juvenile defences against a simulated predator attack (Figures 3B and 3E), exhibited the same delay to return to their eggs after egg defence (Figure 3C) and showed comparable exploration rates when searching for their eggs and juveniles (Figures 3D and 3F).

**Figure 3.**
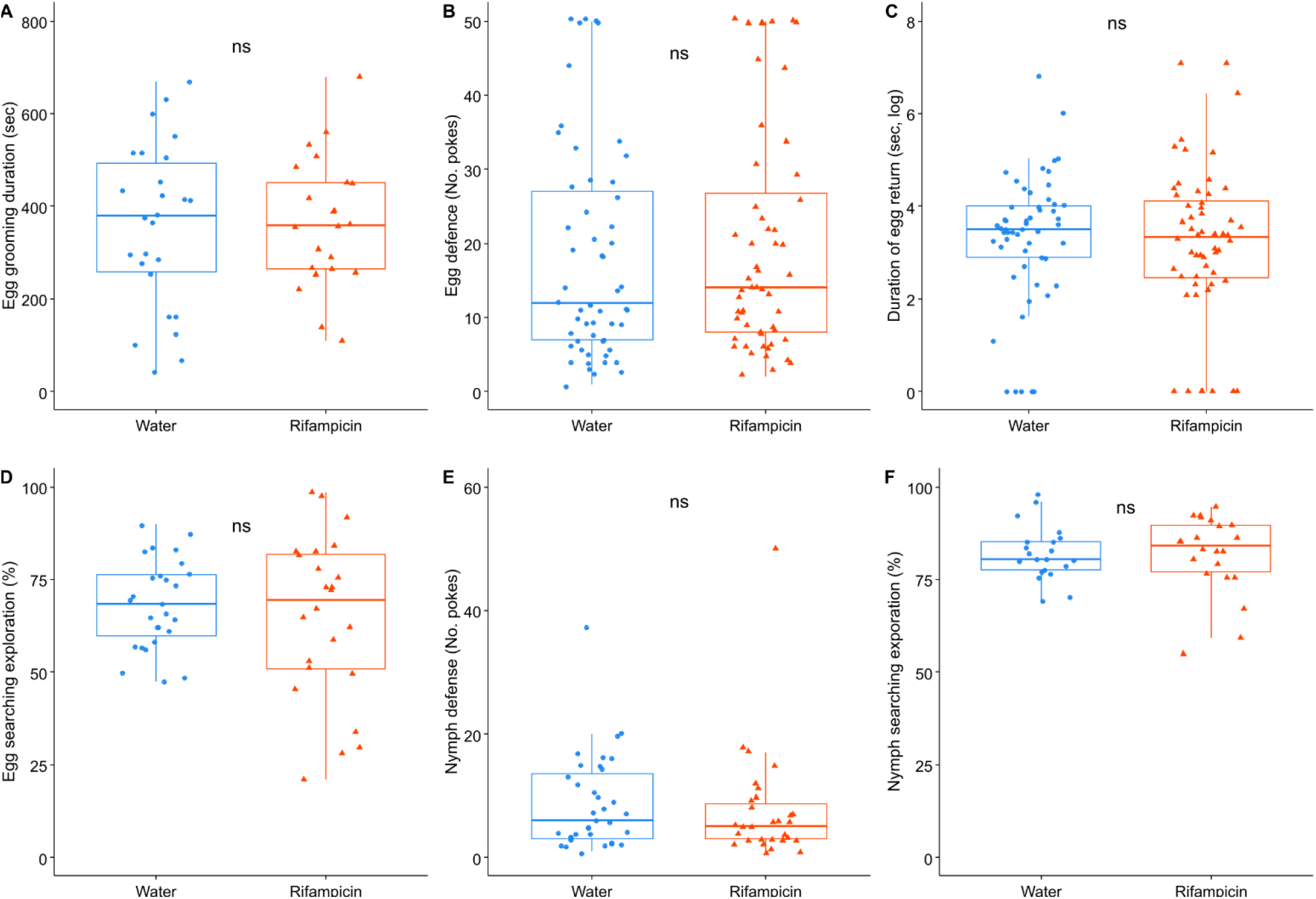
Effect of rifampicin on maternal care. (A) duration of egg grooming, (B) egg defence against a simulated predator, (C) delay of maternal return after egg defence, (D) egg searching, (E) nymph defence against a simulated predator and (F) nymph searching. Box plots depict median (middle bar) and interquartile range (light bar), with whiskers extending to 1.5 times the interquartile range and dots representing experimental values. ns stands for P < 0.05.

### Rifampicin and female’s physiology, reproduction, and longevity

The consumption of rifampicin altered only 3 of the 24 measured proxies of female physiology, reproduction, and longevity. In particular, females fed with rifampicin produced on average twice as many feces pellets per 24h (Figure 4; Table 1), had newly hatched nymphs that were 15% lighter (W = 1244, P = 0.002, adjusted-P = 0.025; Table S1) and laid 2nd clutch eggs that were 7% lighter (W = 628, P = 0.002, adjusted-P = 0.025; Table S1) compared to control females. By contrast, we did not detect any effect of rifampicin on the 21 other traits (Tables 1 and S1).

**Figure 4.**
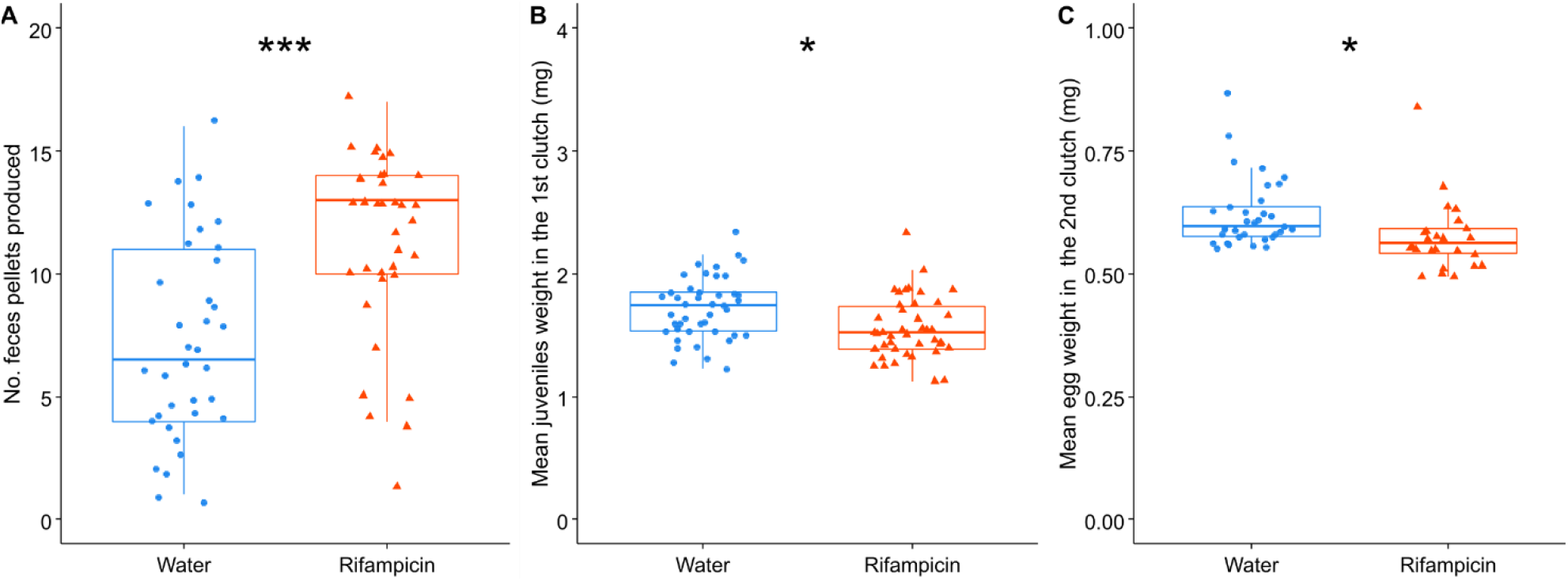
Effects of rifampicin on (A) females’ feces production, (B) mean juveniles weight in the 1st clutch and (C) mean egg weight in the 2nd clutch. Box plots depict median (middle bar) and interquartile range (light bar), with whiskers extending to 1.5 times the interquartile range and dots representing experimental values. ***P < 0.001 and *P < 0.05.

## Discussion

Whereas gut microbial communities shape the physiology, reproduction and behaviour of a great diversity of hosts, their importance on parental care – a key behaviour in social evolution [21–23] - remains poorly explored [30]. In this study, we addressed this gap in knowledge by treating females of the European earwig with rifampicin and measuring the effects on gut microbial communities, maternal care and female physiology, reproduction, and longevity. Our results first reveal that rifampicin altered the composition of the gut microbial community of earwig females and show that this modification diminishes during the period of egg care. Contrary to our predictions, however, the rifampicin-induced alterations of gut microbial communities did not impair the expression of pre- or post-hatching care: rifampicin-treated and control mothers showed similar levels of egg grooming, clutch defences against a predator, maternal return and clutch searching. Independent of maternal care, our results reveal that the consumption of rifampicin increased the females’ production of feces pellets, as well as lead to the production of lighter nymphs and lighter 2nd clutch eggs. By contrast, rifampicin affected none of the other 21 physiological, reproductive and longevity traits measured over the females’ lifetime.

Our experiment first demonstrates that the ingestion of rifampicin by earwig females modified the composition (β-diversity) but not the richness (α-diversity) of bacterial OTUs present in the gut. The fact that rifampicin only shapes indices of β-diversity controlling for phylogeny suggests that OTUs’ phylogeny is the most prominent difference between the community structures present in treated versus non-treated individuals. Because weighted uniFrac is significant and (unweighted) uniFrac is not, our results then indicate that this phylogenetic difference is specific to clades that diverged in the more distant past compared to recently evolved nodes, a pattern in line with broad-spectrum antibiotics acting on conserved bacterial traits. Overall, our findings thus confirm that our treatment successfully altered gut microbial communities in earwigs (just like in other animal species [52–54]). It also indicates that both α- and β-diversity change from pre-oviposition to egg hatching. This stage-specific pattern may result from the absence of food intake for about four weeks before gut sampling in females at egg hatching compared to before oviposition [65], and/or from the different rearing temperatures [66] and differences in female age [67] between the two life stages.

Although gut microbial communities shape the expression of host’s sociality in numerous vertebrate and arthropod species [15–17,19,20], our findings reveal that rifampicin-induced alterations of this community did not affect the expression of pre- and post-hatching maternal care in earwigs. One might have expected that gut microbes directly drive the expression of parental care, as enforcing this social behaviour may allow (at least some) symbionts to reach new hosts (i.e. offspring) that are typically young (thus offering long-lived habitats), display poor immune defences (thus facilitating bacterial establishment and development [68]) and harbour only a few resident microbes (thus limiting the risk of competition within the microbiome [28]). However, our results are at odds with this prediction. This may first suggest that earwig parental care is primarily shaped by microbes that are non-sensitive to rifampicin or that non-sensitive microbes can take over this function (functional redundancy). In insects, gut microbial communities do not only encompass a broad diversity of bacteria (among which some are resistant to rifampicin) but also fungi, protists and other microorganisms that could have key roles and functional redundancies in hosts biology [37,60]. Even if the previous experimental studies linking gut microbiota and host sociality focused on bacteria [15–17,19,20], future studies will be required to confirm that no other members of the gut microbiota shape parental care in our study species, and to explore causal links between the presence of certain members of the microbiome and the level of maternal care expressed by its host. A second potential explanation of our results is that microbial symbionts never developed any specific capabilities to manipulate host sociality, either because adapted strains never occurred within the microbial populations associated with these earwig species (or populations), or because certain antagonistic interactions (e.g. competition) among the members of the microbial community have prevented the emergence of host social manipulation. Any symbiont species (or strain) investing its resources to manipulate host behaviour could indeed be outcompeted within the microbiome by other species or variants that, instead, direct their resources into growth, survival or directly transmission [31](but see [30]). Finally, a third potential explanation is that the symbionts’ capability to manipulate host sociality may change during host social evolution and could thus have vanished in the European earwig. The evolutionary drivers of family life are indeed known to change over time [23] and, while gut microbes may have (at least partly) driven the ancestral evolutionary shift from solitary to family living for the reasons detailed above, the resulting benefits of parental care for the hosts could have consolidated the expression of care and thus reduced the capability of symbionts to control host social behaviours and/or reduced the sensitivity of the hosts to this manipulation. Based on this hypothesis, alterations in gut microbial communities should not shape the expression of parental care once this behaviour is established. This process could have limited the maintenance of symbiont control over parental care in earwigs, where maternal care is well established. Notwithstanding the drivers of our results, our findings provide first experimental evidence that broad alteration of the gut microbiota (with rifampicin) does not directly or indirectly impair the expression of maternal care, and thus call for caution when considering the role of gut microbiota in this important social behaviour.

Despite its apparent lack of effects on maternal care, rifampicin altered three maternal traits related to physiology and reproduction. The first trait is the production of feces pellets, which was twice as high in rifampicin compared to control females. This result was not surprising, as the gut microbiota often plays a key role in nutrient extraction and digestion [69] and its alteration by antibiotics typically disturbs the host’s digestive efficiency and triggers an overproduction of fecal material. The two other traits were the weights of the 2nd clutch juveniles and 2nd clutch eggs, which were (slightly) lighter in rifampicin compared to control females. Light eggs and newly hatched juveniles are often thought to reflect low offspring quality in insects [70], and further studies are required to confirm this association in earwigs.

Rifampicin altered none of the 21 others physiological, reproductive and longevity traits measured in earwig mothers. Whereas these findings contrast with a large body of literature showing the broad impact of altered gut microbial communities on host biology [4], they are in line with a few recent studies showing that antibiotic-induced alteration of gut microbial communities does not affect the development and survival of three Lepidoptera species (Danaus chrysippus, Ariadne merione and Choristoneura fumiferana [71–73]). Together with these findings, our results thus provide support to the idea that essential microbial symbioses are not universal across insect species [71,74]. In these lepidopterans, the lack of microbial symbioses has been explained by the fact that they do not depend on specific gut bacteria to derive critical nutrition from their dietary resources [73,75]. This might also be the case in the European earwig because it is omnivorous [42] and thus a partnership with bacteria facilitating the digestion of specific food sources might not have been required during species evolution. Follow-up studies will investigate whether (and which part of) the earwigs gut microbiota is transient.

To conclude, our study reveals that rifampicin consumption alters female gut microbial communities in earwigs, but provides no evidence for a link between this alteration and the expression of maternal care, as well as no evidence for a strong impact of this alteration on earwig physiology, reproduction and survival. Overall, these findings provide support to a recent proposal that microbial enforcement of host social interactions is unlikely to evolve [31] and to the emerging idea that not all animals have evolved a co-dependence with their microbiome [71,74]. Nevertheless, shedding light on whether and how a symbiotic community shape hosts biology is a difficult task, mostly due to the number of players possibly involved and the complexity of their potential interactions [72]. Hence, our findings call for follow‐up studies testing whether and how other members (non-sensitive to rifampicin) of the gut microbial community could shape the expression of parental care in family-living animals and/or drive important fitness parameters of earwig biology.

## Data accessibility

The raw metabarcoding sequence data have been deposited in the NCBI Sequence Read Archive under the BioProject PRJNA646389 with BioSample accession numbers SAMN15547835 to SAMN15547872. The Dataset and R script used for analyses of life-history traits and behaviour as well as the detailed bioinformatics pipelines reported in this manuscript are available on Zenodo [76].

## Supplementary material

Script and codes are available online: https://doi.org/10.5281/zenodo.4506829

Supplementary materials are available online: https://doi.org/10.1101/2020.10.08.331363

## Ethics approval

No animal ethics approval was required to conduct this study. All individuals were nevertheless handled with care following the 3Rs (Replacement, Reduction and Refinement) for the ethical use of animals in testing.

## Acknowledgements

We thank Jos Kramer, Maximilian Körner, as well as Nadia Aubin-Horth, Gabrielle Davinson, an anonymous reviewer and Trine Bilde (Peer Community in Evolutionary Biology referees and recommender, respectively) for their insightful comments on this manuscript. This research has been supported by a research grant from the French Ministry of Research (to S.V.M.) and a research grant from the Agence Nationale de la Recherche (ANR-20-CE02-0002 to J.M.). Version 5 of this preprint has been peer-reviewed and recommended by Peer Community In Evolutionary Biology (https://doi.org/10.24072/pci.evolbiol.100121)

## Appendix

Appendix 1 and Table S1, S2 and S3

