## Supplementary material for "Alteration of gut microbiota with a broad-spectrum antibiotic does not impair maternal care in the European earwig"

### Figure S1 – Timeline of the experiment.

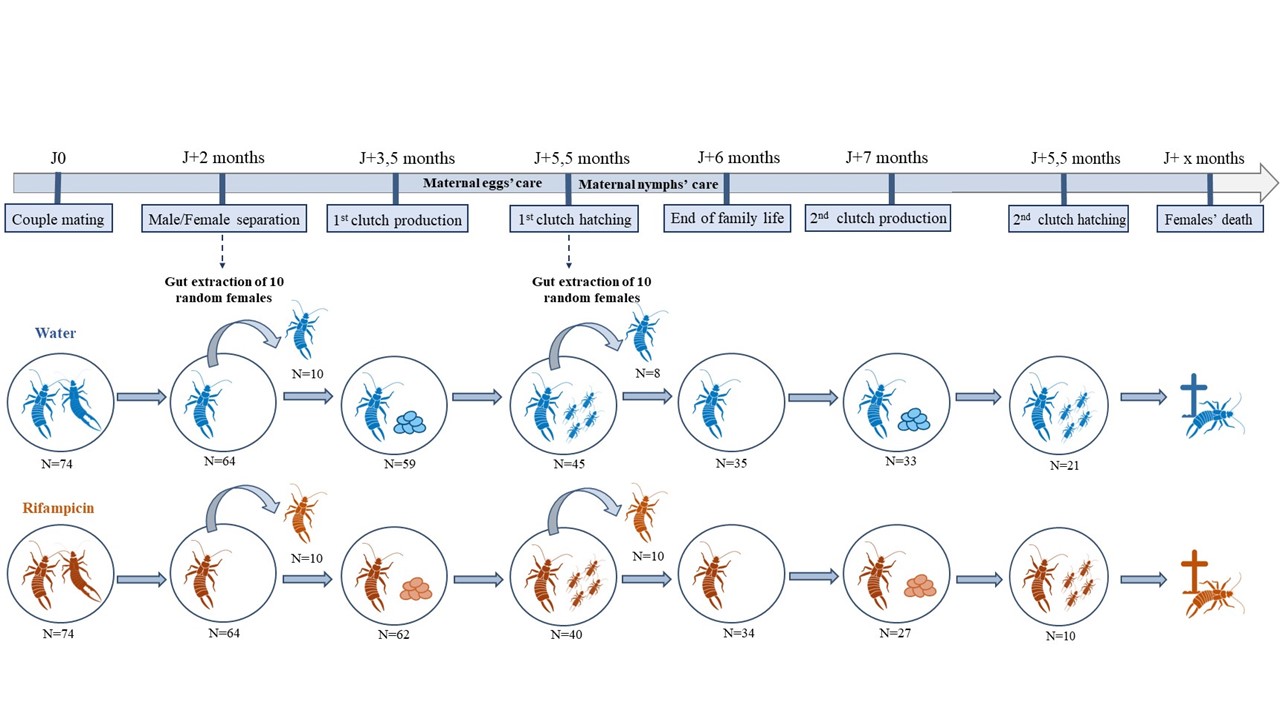

### Blue and red individuals received the water (control) and antibiotic treatments, respectively. N indicates the total number of replicates used at each step.

### DNA extraction and PCR

Genomic DNA was extracted from guts using Nucleo-Spin Tissue extraction kit, following the manufacturer instructions. Before the lysis step, a sterile stainless-steel bead (5 mm, Qiagen) was placed in each tube and samples were ground using a Tissue-Lyser II (Qiagen) for 30 s at 30 Hz.

For each PCR (Polymerase Chain Reaction), 3.5 µl of MTP Taq buffer 10× (Merck), 0.7 µl of dNTP mix (10 mM, Merck), 0.88 µl of each primer (20 µM), 0.6 µl of MTP Taq DNA Polymerase (Merck) and 25.45 µl of water (Merck) were mixed with 3 µl of DNA template. PCR amplifications were performed with 1 min at 94°C followed by 30 cycles composed of 1 min at 94°C, 1 min at 65°C and 1 min at 72°C, and a final step at 72°C for 10 min.

### Bioinformatics pipeline

At the GeT-PlaGe genomic platform, the quality of the run was checked using PhiX (Illumina), each pair-end sequence was assigned to its sample and pair-end reads were assembled using Flash software [1]. The sequencing reads were processed following the FROGS standard operation procedure [2]. A pre-process tool was used to remove sequences without primers, with not expected lengths, containing ambiguous bases, and to trim the primers. Sequence clustering was carried out with SWARM [3] with a first denoising step (aggregation distance = 1) followed by a second clustering step (aggregation distance = 3). The aggregation distance corresponds to the maximum number of differences between sequences in each aggregation swarm step. The denoising step allows to build very fine clusters with minimal differences, and the second clustering step was run between seeds from the first aggregation step. Chimeric sequences were detected and removed with VSEARCH [4], and a filtering tool was used to remove clusters with a read number abundance of less than 0.005 % of all sequences [5]. Sequences were finally classified with SILVA taxonomy version 138 [6,7] for V3V4 reads.

### Negative controls through library preparation and sequencing

Several controls have been carried out during the experiments, and as expected, all these controls were negative. Therefore, no library preparation or sequencing was produced. First, we checked the absence of inter-sample contamination by taking samples from the dissection tools (forceps) and the T1 buffer. Then, we extracted the DNA from these controls and verified the absence of 16S amplification by PCR. During this same amplification phase, we also placed 2 negative controls (water as template) to verify the absence of contamination during the PCR. At the sequencing platform (i.e., the GeT-PlaGe platform), and prior to sequencing, the lack of contamination was checked with a negative control during the second PCR step (water as template). Again, all these controls were as negative.
